# Genetic and immune landscape evolution defines subtypes of MMR deficient colorectal cancer

**DOI:** 10.1101/2022.02.16.479224

**Authors:** Benjamin R. Challoner, Andrew Woolston, David Lau, Marta Buzzetti, Caroline Fong, Louise J. Barber, Gayathri Anandappa, Richard Crux, Ioannis Assiotis, Kerry Fenwick, Ruwaida Begum, Dipa Begum, Tom Lund, Nanna Sivamanoharan, Harold B. Sansano, Melissa Domingo-Arada, Amina Tran, Bryony Eccles, Richard Ellis, Stephen Falk, Mark Hill, Daniel Krell, Nirupa Murugaesu, Luke Nolan, Vanessa Potter, Mark Saunders, Kai-Keen Shiu, Sebastian Guettler, James L. Alexander, Héctor Lázare-Iglesias, James Kinross, Jamie Murphy, Katharina von Loga, David Cunningham, Ian Chau, Naureen Starling, Juan Ruiz-Bañobre, Tony Dhillon, Marco Gerlinger

**Author notes:** **Corresponding Author:** Dr Marco Gerlinger, Barts Cancer Institute, Queen Mary University of London, UK. Gastrointestinal Cancer Unit, The Royal Marsden NHS Foundation Trust, London, UK. The Institute of Cancer Research, London, UK.

## Abstract

Mismatch repair deficient colorectal cancers have high mutation loads and many respond to immune checkpoint-inhibitors. We investigated how genetic and immune landscapes co-evolve in these tumors. All cases had high truncal mutation loads. Driver aberrations showed a clear hierarchy despite pervasive intratumor heterogeneity: Those in WNT/βCatenin, mitogen-activated protein kinase and TGFβ receptor family genes were almost always truncal. Immune evasion drivers were predominantly subclonal and showed parallel evolution. Pan-tumor evolution, subclonal evolution, and evolutionary stasis of genetic immune evasion drivers defined three MMRd CRC subtypes with distinct T-cell infiltrates. These immune evasion drivers have been implicated in checkpoint-inhibitor resistance. Clonality and subtype assessments are hence critical for predictive immunotherapy biomarker development. Cancer cell PD-L1 expression was conditional on loss of the intestinal homeobox transcription factor CDX2. This explains infrequent PD-L1 expression by cancer cells and likely contributes to the high recurrence risk of MMRd CRCs with impaired CDX2 expression.

## INTRODUCTION

DNA mismatch repair deficient (MMRd) colorectal cancers (CRCs) are molecularly and clinically distinct from MMR proficient (MMRp) CRCs (Germano et al., 2018). Loss of MMR protein expression (MLH1, PMS2, MSH2 or MSH6) in MMRd CRCs establishes a hypermutator phenotype and high mutation loads. The large number of mutation-encoded neoantigens, as well as activation of the cGAS-STING pathway in cases with MLH1 loss (Guan et al., 2021; Lu et al., 2021), make these cancers highly immunogenic. Following surgical resection, stage 1/2 MMRd CRCs have a lower recurrence risk compared to stage-matched MMRp CRCs, likely through better control by the immune system. However, the favorable prognostic impact of MMRd is lost in tumors with lymph node metastases (stage 3) (Domingo et al., 2016). To what extent the evolution of immune evasion (IE) mechanisms contributes to the transition from localized to metastatic disease remains unclear (Kloor et al., 2010). Furthermore, immunotherapy with PD1/PD-L1 checkpoint-inhibitors (CPIs) is ineffective in metastatic (stage 4) MMRp CRCs (Le et al., 2015) but has high response rates in MMRd CRCs (43.8%) (Andre et al., 2020), confirming an important role for the PD1/PD-L1 immune-checkpoint in IE. An early-phase trial showed even higher response rates of 69% with combined PD1 and CTLA4 CPIs in metastatic disease (Lenz et al., 2020) and of 100% in early-stage MMRd CRCs with 12 out of 19 tumours (63%) achieving a pathological complete response (Chalabi et al., 2020). However, combination immunotherapy is significantly more toxic. Thus, there is a major need to identify biomarkers to predict who benefits from single-agent CPI, who needs combination CPIs and to develop better therapies for those that do not respond to either approach.

Biomarkers for CPIs in other tumor types include mutation loads, the number of insertions and deletions (indels) (Turajlic et al., 2017), as well as cytotoxic CD8 T-cell infiltrates and expression of PD-L1 (Havel et al., 2019). Neither of these correlated with immune infiltrates or CPI responses in MMRd CRC (Maby et al., 2015; Overman et al., 2017). Moreover, Inactivation of genes in the IFNγ signal transduction pathway or of important components of MHC class I antigen presentation, such as *B2M*, have been associated with resistance to CPI therapies in several tumour types (Gao et al., 2016; Sade-Feldman et al., 2017; Shin et al., 2017; Zaretsky et al., 2016). Data in MMRd CRCs are less clear as low expression of B2M was not associated with CPI resistance and murine tumours with B2M loss still responded to combined PD1 and CTLA4 inhibition (Germano et al., 2021). Importantly, the hypermutator phenotype distinguishes MMRd tumors from other CPI-sensitive cancers such as melanoma, lung or renal cancer which have low mutation rates during progression. We have shown in MMRd gastro-esophageal cancers that it promotes extreme intratumor heterogeneity (ITH) (von Loga et al., 2020). Although this complicates biomarker identification (Gerlinger et al., 2014; Gulati et al., 2014), ITH has not been systematically assessed in MMRd CRCs. Driver evolution can follow recurrent patterns (de Bruin et al., 2014; Gerlinger et al., 2014), suggesting some order despite evolution which is fuelled by randomly generated genetic aberrations. Biomarker development hence requires definition of which drivers are commonly truncal and can be assessed reliably from single samples, and which require assays competent to address ITH. Beyond clarifying suitable biomarker strategies to assess who should be treated with PD1 inhibition and who requires combined PD1 and CTLA4 therapy, interrogating subclonal driver evolution may furthermore provide a better understanding of progression to lethal disease and the identification of prognostic biomarkers. Here, we apply multi-region multi-omics and immunohistochemistry (IHC) analysis to reveal ITH of driver aberrations and immune infiltrates and how they co-evolve in MMRd CRCs.

## RESULTS

20 CRCs where immunohistochemistry (IHC) reported MMRd were analysed consecutively (**Table 1**). 55 primary tumor regions, 15 lymph nodes and one distant metastasis were sequenced with a panel of 194 genes which are recurrently mutated in MMRd or MMRp CRCs, or implicated in immune evasion and may confer CPI resistance (Supplementary table 1) (Cortes-Ciriano et al., 2017; Gao et al., 2016; Giannakis et al., 2016; Grasso et al., 2018; von Loga et al., 2020; Zaretsky et al., 2016). Targeted sequencing enabled high sequencing depths (median: 2659x) which is critical to avoid ITH overestimation in samples with low cancer cell content such as lymph node metastases or tumor regions with high immune cell infiltrates.

**Table 1:** Patient and pathological characteristics.

|  |  |
| --- | --- |
|  | MMRd CRC Cohort<br>(n=20) |
| **Median age at resection (range)** | 61.2 (33.5-79.2) |
| **Sex** |  |
| Male | 30% (6) |
| Female | 70% (14) |
| **Stage (AJCC/UICC 8th edition)** |  |
| 1 | 15% (3) |
| 2 | 15% (3) |
| 3 | 65% (13) |
| 4 | 5% (1) |
| **Predominant Differentiation** |  |
| Well to moderate | 45% (9) |
| Poor | 25% (5) |
| Mucinous | 30% (6) |

The median non-silent mutation load of individual tumor regions was 58 (range: 10-90, **Fig.1A**, see Supplementary table 2 for detailed patient characteristics and sequencing results). Using a linear regression model trained on 518 CRC exomes from the Cancer Genome Atlas (Bailey et al., 2018) (Supplementary figure 1) this extrapolated to a median of 2418 mutations across the whole exome in our series. 19 tumors had high mutation loads (28-90 mutations/region, estimated 1102-3822/exome) that were consistent with expectations for MMRd. T20 was an outlier with only 10 mutations (estimated 313/exome), suggesting this was an MMRp tumor. Histology review showed patchy rather than complete loss of MLH1/PMS2 which is a recognised feature that can cause misclassification as MMRd (Markow et al., 2017). This tumor was excluded from the analysis. A median of 44 mutations (range: 20-70) were ubiquitously present in all regions of individual tumors (extrapolated 1844 mutations/exome, range: 752-2945). Thus, all MMRd CRCs had high truncal mutation loads.

**Figure 1:**
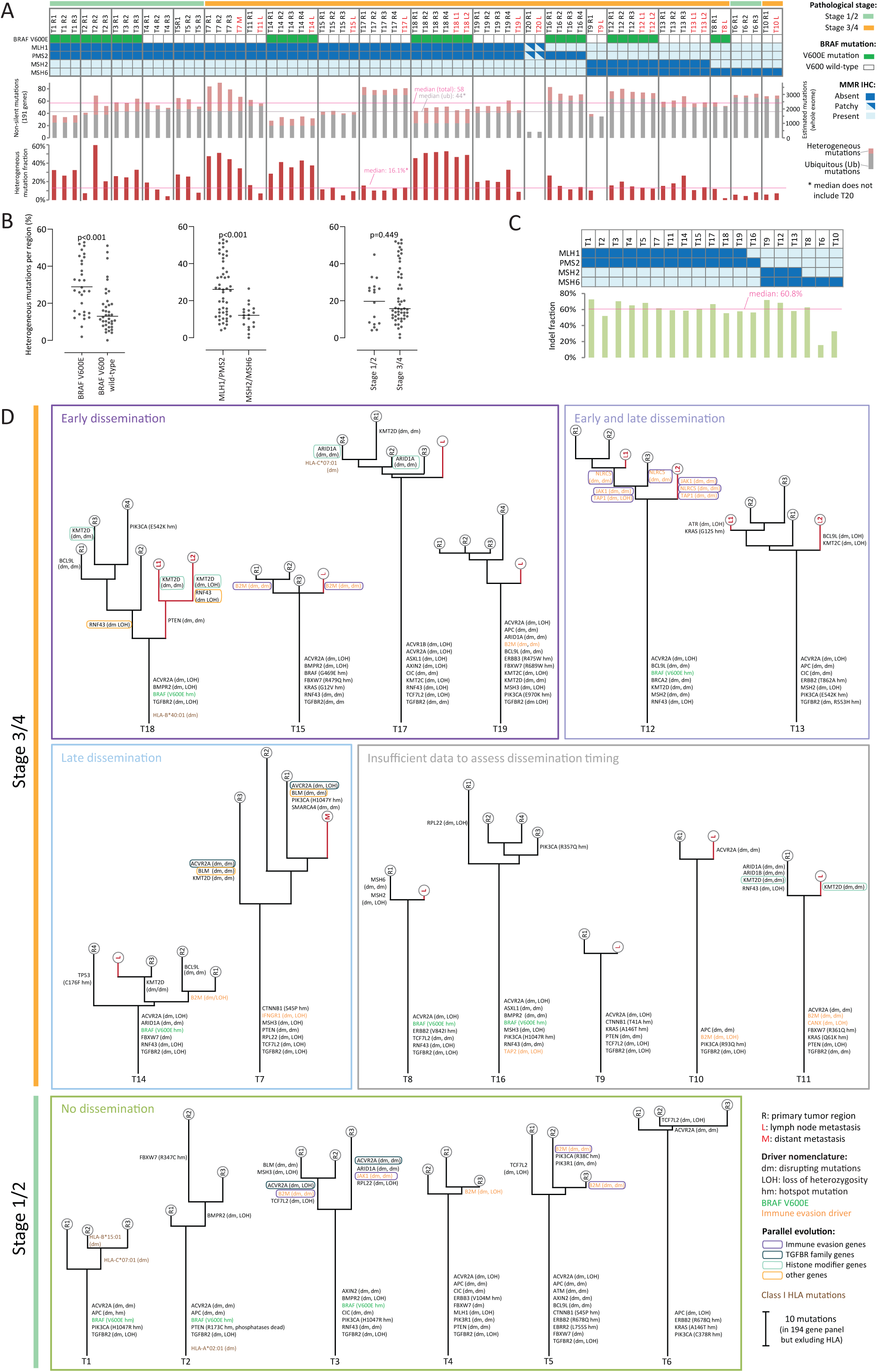
Multi-region sequencing analysis of MMRd CRCs. A. Ubiquitous and heterogeneous mutations detected in 191 of 194 genes (excluding HLA-A/B/C) by tumor region/metastasis. Stage, *BRAF* V600E mutations and MMR protein expression are shown. The y-axis on the right shows the estimated mutation load in the exome which was extrapolated by linear regression. B. Fraction of heterogeneous mutations per region by BRAF mutation status, MMR protein loss and stage. The horizontal lines represent the median. Significance was assessed with the Mann-Whitney test. C. Fraction of indels among all called mutations by MMR protein loss. D: Phylogenetic trees generated from mutation calls with the Phylip parsimony algorithm Pars, grouped according to stage and metastatic dissemination timing. Driver aberrations in tumor suppressor genes or immune evasion genes were mapped on the branch where the second disrupting aberration had been acquired and in oncogenes on the branch where a hotspot mutation had been acquired. T: tumor, R: primary tumor region, L: lymph node metastasis, M: distant metastasis.

### Intratumor heterogeneity and indels by subtype and stage

ITH was identified in all cases (median: 16.1% heterogeneous mutations/region, **Fig.1A**). It was significantly higher in *BRAF* V600E (median 26.9%) versus V600 wild-type tumors (13.0%) and in tumors with MLH1/PMS2 loss (25.7%) compared to those with MSH2/MSH6 loss (12.1%, **Fig.1B**). ITH was similar in stage 1/2 and stage 3/4 tumors. Ubiquitous mutation loads did not significantly differ by *BRAF* mutation, MMR pattern or stage (Supplementary figure 2).

**Figure 2:**
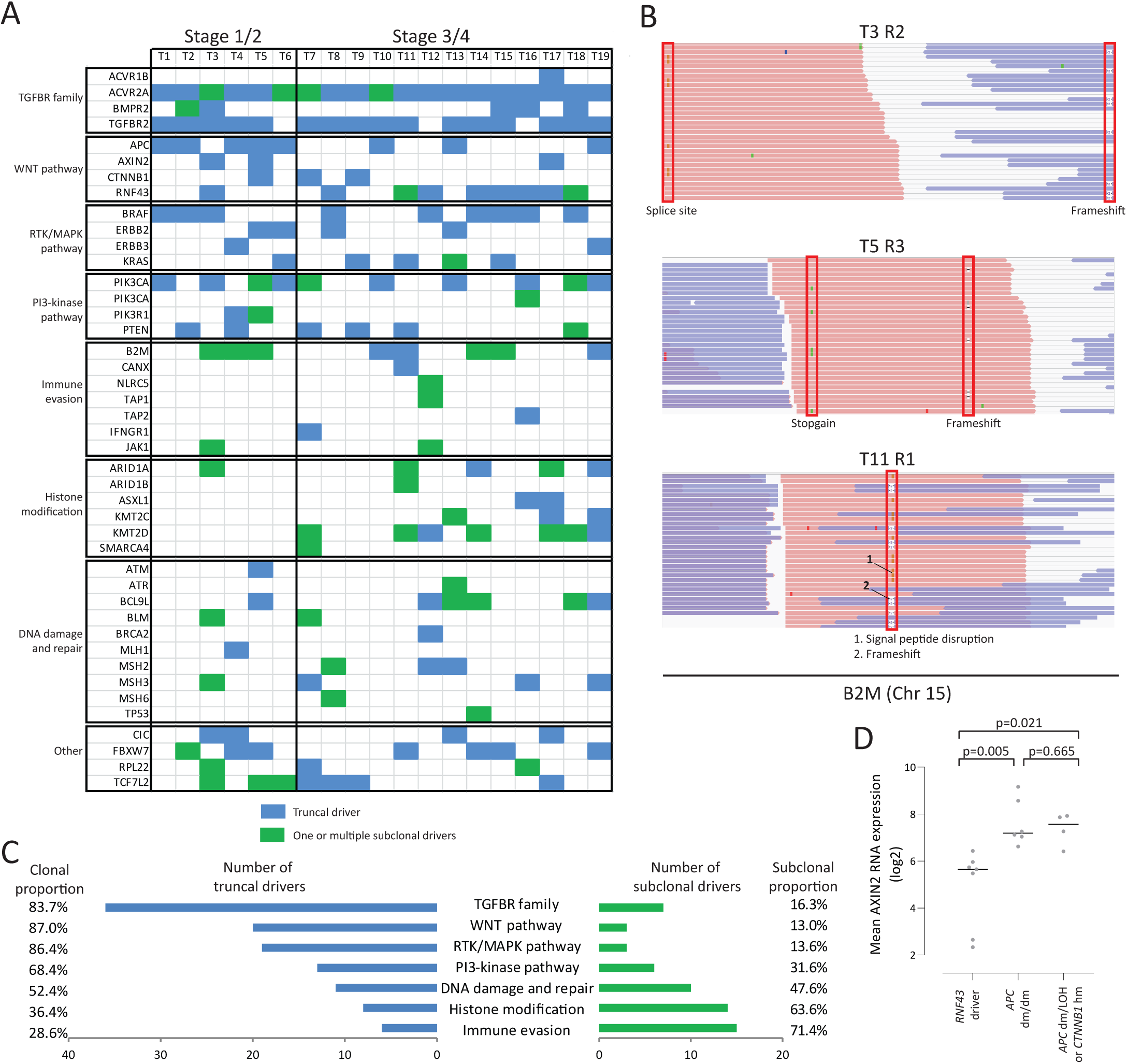
Driver aberrations and heterogeneity. A. Heatmap of identified truncal or subclonal driver aberrations for each tumor. B. Confirmation that distinct *B2M* mutations were located on different paired-end reads but never together. Screenshots were obtained from the Integrated Genomics Viewer. C. Proportion of truncal and subclonal driver aberrations by functional group. D. Mean AXIN2 expression in primary tumor regions by truncal driver aberrations in *APC, RNF43* and *CTNNB1*. Horizontal lines represent the mean and significance was assessed with the t-test.

A high proportion of indels is characteristic of MMRd and a median of 60.8% of non-silent mutations were indels (**Fig.1C**). The higher proportion of indels compared to previous reports in MMRd tumors (Turajlic et al., 2017) is likely due to the overrepresentation of microsatellite repeats among driver genes in our sequencing panel. Two tumors had noticeably lower indel fractions (T6: 15.6%, T10: 31.5%) and both only showed isolated MSH6 loss. This can be explained by the predominant recognition of base-base mismatches by the MSH2/MSH6 heterodimer, whereas the MSH2/MSH3 and MLH1/PMS2 heterodimers are also important for recognition and repair of indels (Houlleberghs et al., 2017). A third tumor with isolated MSH6 loss had a high indel fraction (T8). Despite preserved MSH2 protein expression, this tumor harbored an early somatic stop-codon mutation in *MSH2* (E28X) in all regions and loss of heterozygosity in R1. Lynch syndrome patients with *MHS2* start-codon mutations and preserved MSH2 expression have been described, suggestive of a hypomorphic MSH2 isoform expressed from a downstream start codon (M67 in the canonical transcript) (Kets et al., 2009). Expression of this alternative isoform likely explains the high indel fraction despite preserved MSH2 protein expression in T8.

### Metastasis timing revealed by phylogenetic analysis

We next generated phylogenetic trees to assess how tumors had evolved (**Fig.1D**). All tumors showed branched evolution. Most strikingly, metastases diverged before detectable genetic diversification of the primary tumor in 6/8 cases (75%) where multiple primary tumor regions and at least one metastatic site were available for dissemination timing assessment. The ability to metastasize was hence likely acquired on the phylogenetic trunk, suggesting that most stage 3/4 cancers were already ‘born bad’ (Sottoriva et al., 2015).

### Driver aberration evolution is characterized by a clear hierarchy

The acquisition of new genetic drivers is arguably the most relevant consequence of cancer evolution. We identified drivers by cataloguing hotspot mutations in known oncogenes as well as disrupting aberrations (frameshift, splice-site, nonsense mutations, disruption of the signal peptide, and loss of heterozygosity) that were likely to confer biallelic inactivation of tumor suppressor (TS) genes or immune evasion (IE) driver genes (**Fig.2A**, driver genes assessed: Supplementary table 3, see methods for the algorithm used to identify likely drivers). A hotspot mutation (hm) in an oncogene or loss of heterozygosity (LOH) of a TS or IE gene combined with a disrupting mutation (dm) that is clonal in a sample reliably defines driver aberrations. Yet, many TS and IE genes harbored two dm in these hypermutated tumors. Even when clonal, two distinct dm can only inactivate a gene if they affect both alleles. Most detected dm were not in close enough proximity to assess in the short-read sequencing data whether they were present on different alleles or on the same allele. However, of 5 cases with multiple *B2M* mutations in the same sample, 3 harbored dm pairs within the span of paired-end sequencing reads. In all three, mutations were mutually exclusive, i.e. never detected together in a single read (**Fig.2B**). We therefore accepted two independent clonal dm in a TS or IE gene as a marker of likely biallelic inactivation. All identified driver aberrations (**Fig.2A**, Supplementary table 4) were mapped onto the phylogenetic trees (**Fig.1D**).

Analysis of truncal and subclonal driver acquisition showed a clear hierarchy of driver evolution (**Fig.2C**). WNT/βCatenin pathway (WNT) aberrations, activating mutations of receptor tyrosine kinases/mitogen activated protein kinase pathway (RTK-MAPK) and inactivation of TGFβ-receptor family members (TGFBR) were almost always truncal (87.0%, 86.4% and 83.7%, respectively). Moreover, these pathways were altered on the trunk of 89.5%, 94.7% and 84.2% of tumors, respectively, demonstrating that they are critical for tumor initiation. Mutual exclusivity was observed for disrupting *APC* and *RNF43* aberrations in the WNT-pathway. This allowed further validation of our algorithm for the identification of drivers in TS genes as AXIN2 RNA overexpression is an established marker of ligand-independent WNT activation (described to occur through *APC* inactivation or *CTNNB1* hm) versus ligand dependent WNT activation (through *RNF43* inactivation) (Kleeman and Leedham, 2020). Six tumors that harbored two truncal dm in *APC* showed significantly higher AXIN2 expression than tumors with truncal *RNF43* drivers (**Fig.2D**), and similar expression to those with truncal *APC* dm/LOH or *CTNNB1* hm. Thus, the detection of two independent dm, even in a large TS gene such as *APC*, strongly indicates biallelic inactivation.

Aberrations activating the PI3-kinase pathway (*PIK3CA* hm, inactivation of *PTEN* or of *PIK3R1*), inactivation of histone modifiers or of DNA damage response and repair (DDR) genes were truncal in 36.4%-68.4% of occurrences. Inactivation of HLA class I antigen presentation or interferon gamma (IFNγ) signalling genes, previously shown to enable IE and resistance to checkpoint-inhibitor treatment (Gao et al., 2016; Sade-Feldman et al., 2017; Shin et al., 2017; Zaretsky et al., 2016), were predominantly subclonal (71.4%). Thus, ITH impedes accurate assessment of these critical immunotherapy biomarkers.

Disrupting mutation of individual HLA class I genes were infrequent (5 mutations in T1, T2, T18, T17; **Fig.1D**) in comparison to drivers in other antigen processing/presentation genes (16 mutations). This suggests that disruption of genes with broad effects on antigen presentation is a more effective evolutionary path to immune evasion in tumors with such high neoantigen numbers.

### Parallel evolution predominates among IE driver genes

We next assessed parallel evolution which is a strong indicator of Darwinian selection (Gerlinger et al., 2014; Gerlinger et al., 2012). This was defined by the presence of at least two distinct driver aberrations affecting genes with similar function on different phylogenetic branches. 8/19 tumors (42.1%) showed 12 instances of parallel evolution in 9 different driver genes (**Fig.1D**). Parallel evolution was commonest for IE genes (6/9 genes), most frequently affecting *B2M*. Parallel evolution of *KMT2D* and *ACVR2A* inactivation was present in two tumors, each, and *BLM, ARID1A* and *RNF43* in one. The most striking example was observed in T12 where *NLRC5* (the master regulator of HLA class I expression) was inactivated in three and the antigen transporter *TAP1* and the IFNγ-pathway signalling gene *JAK1* each in two different ways, leading to pan-tumor inactivation of these three genes in all regions. Parallel evolution of IE occurred in 4/6 tumors that harbored any subclonal IE drivers, indicating that these tumors were under intense immune selection pressure. Subclonal IE drivers were absent in all 5 tumors with truncal IE drivers. Parallel evolution, and truncal and subclonal mutual exclusivity of IE drivers in a total of 11/19 tumors (57.9%) substantiates that these evolved through Darwinian selection.

### Driver heterogeneity between primary tumors and metastases

We questioned whether specific drivers predominantly evolved in the 15 metastases (**Fig.1D**). Only *KMT2D* acquired driver aberrations more than once (3/15 metastases). IE drivers were acquired by 2 metastases; *B2M* in T15 and *JAK1, NLRC5* as well as *TAP1* in T12. *PTEN* was inactivated on the common branch of two metastases in T18. No new drivers had evolved on the branches leading to 7/15 metastases. Thus, drivers can be unique to metastatic sites but we found no strong evidence for recurrent selection of specific drivers, consistent with our above observation that metastatic potential may be encoded on the trunk in most cases.

### Heterogeneity of immune infiltrates

T-cell infiltrates are biomarkers of cancer immunogenicity and checkpoint-inhibitor sensitivity (Havel et al., 2019). We assessed these by quantifying CD8 T-cells with IHC and computational image analysis (Supplementary table 2). Mean CD8 T-cell densities in primary tumor regions showed large variability between cases (1.9%-21.5% of nucleated cells), questioning how T-cell density is regulated. It did not significantly differ by *BRAF* status or MMR loss patterns, nor between stage 1/2 and stage 3/4 tumors (**Fig.3A**). Correlating CD8 T-cells in the primary tumor with mutation loads, the percentage of heterogeneous mutations, ubiquitous mutation loads or ubiquitous indel loads showed no significant association (Supplementary table 5). This is perhaps unsurprising as all tumors had sufficient ubiquitous mutations to encode for multiple neoantigens, even when taking conservative estimates that only 0.5-1% of somatic mutations give rise to HLA-presented neoantigens (Newey et al., 2019; Schumacher et al., 2019). However, the realized antigenicity may vary due to the impact of abundant IE drivers.

**Figure 3:**
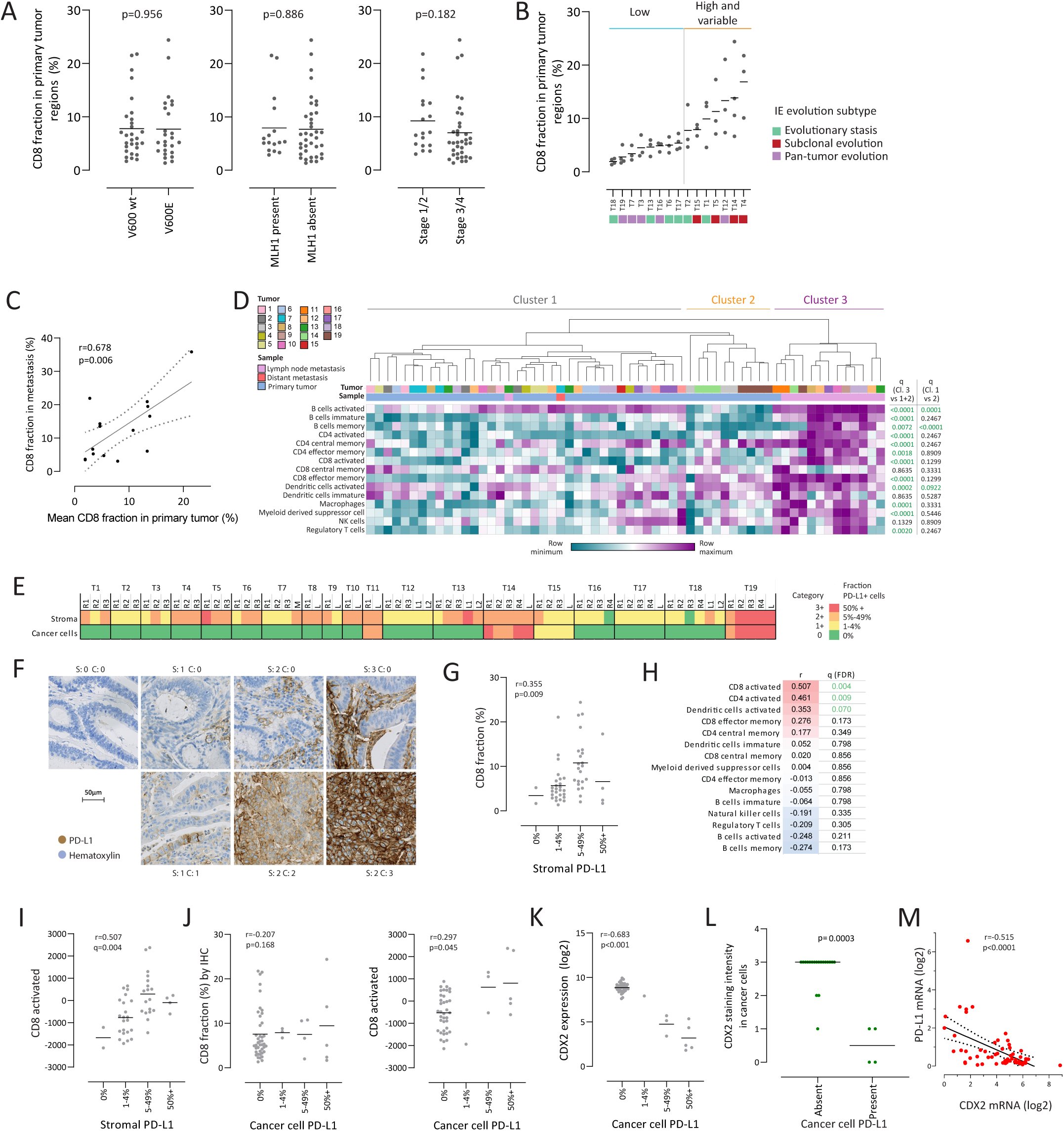
Immune landscapes of MMRd CRCs. A. Fraction of CD8 T-cells (IHC) among all nucleated cells per primary tumor region by *BRAF* mutation status, MLH1 loss and stage. B. CD8 T-cell fraction (IHC) in each primary tumor region and mean for each tumor where at least 3 primary tumor regions were available. C. Linear regression analysis of average CD8 T-cell fraction in the primary tumor and in matched lymph node metastases. The regression line (solid line) and 95% confidence intervals (dotted lines) are shown. D. Euclidian clustering of 15 immune cell subtype abundances inferred from RNA expression data with ssGSEA. q values were generated with Morpheus with the t-test and multiple testing correction. E. Proportion of PD-L1 positive stromal cells and cancer cells by IHC in each analysed sample. F. Examples of PD-L1 staining in stroma and cancer cells (S: stromal staining, C: cancer cell staining). G. CD8 T-cell fraction (IHC) in primary tumor regions by PD-L1 positivity in stromal cells. H. Correlation of immune cell subtype abundance assessed by ssGSEA with stromal PD-L1 expression. I. Individual sample data of activated CD8 T-cell abundance (inferred from RNA expression) in primary tumor regions by PD-L1 positivity. J. CD8 T-cell fraction IHC (left) and activated CD8 T-cells from RNA expression (right) in primary tumor regions by PD-L1 positivity in cancer cells. K. CDX2 RNA expression in primary tumor regions by PD-L1 positivity in cancer cells. L: CDX2 staining intensity (0: absent, 1: weak, 2: moderate, 3: strong) by cancer cell PD-L1 expression in an independent validation cohort of 23 MMRd CRCs. M: Linear regression analysis of PD-L1/CD274 and CDX2 mRNA expression in 57 colorectal cancer cell lines from the Cancer Cell Line Encyclopedia. Horizontal bars show the medians in panel L and means elsewhere. p-values were calculated with the Mann Whitney test (L) and with the t-test for other panels. False discovery rate multiple testing correction was applied where q values are shown. p<0.05 and q<0.1 were considered significant. Correlation coefficients (r) were calculated with Pearson (C, M) and Spearman rank (G-K) tests.

CD8 T-cell densities also varied within primary tumors (**Fig 3B**). Dichotomisation of tumors using the median value across primary tumor region CD8 T-cell fraction means, distinguished tumors with low (mean: 4.0%) from tumors with high, albeit frequently variable, CD8 T-cell infiltrates (mean: 11.7%). This suggested a tumor-intrinsic setpoint which is however accompanied by marked variability in tumors with dense infiltrates. To further test this hypothesis, we correlated CD8 T-cells in the primary tumor versus metastases. T-cell densities were significantly higher in metastases than primary tumors (paired t-test: p=0.028, mean increase: 1.53-fold) but showed a significant correlation (r=0.678, **Fig.3C**). This indicated that T-cell densities are regulated by tumor specific characteristics and also influenced and augmented by the microenvironment in metastases.

To obtain more granular insights, we computationally inferred 15 immune cell subtypes from RNA expression data (Immune cell abundances: Supplementary table 2). IHC-measured CD8 T-cells correlated most strongly with activated CD8 T-cells (r=0.525, q<0.001, Supplementary figure 3), confirming that inference from expression data worked reliably. Hierarchical clustering showed that all metastases except one lymph node and the distant metastasis clustered together (**Fig 3D**). This cluster (cluster 3) was significantly enriched for immune cells expected in the lymph node environment such as B-cells and activated dendritic cells, but also for multiple other immune cell types including activated CD8 and CD4 cells. Primary tumor regions from individual cancers co-segregated into one of the two other major clusters. The only significant differences between these were higher activated and memory B-cells in cluster 1 and enrichment of activated dendritic cells in cluster 2 (**Fig 3D**).

### Distinct mechanisms influence PD-L1 expression by stromal and cancer cells

Expression of PD-L1 by cancer cells, stromal cells or both is a major non-genetic IE mechanism (Juneja et al., 2017; Kleinovink et al., 2017; Lau et al., 2017). We used IHC and categorized samples with accepted cut-offs (0: 0%; 1+: 1-4%; 2+: 5-49%; 3+: 50-100%) based on the percentage of PD-L1 positive cancer or stromal cells (Supplementary table 2) (Moller et al., 2021). Stromal expression was common, with 97.1% of regions showing 1+ to 3+ staining (**Fig.3E-F**). Heterogeneity within tumors was modest. Only 4 tumors harbored PD-L1 expressing cancer cells (23.2% of all regions) and expression was detected in all regions of these. Cancer cell PD-L1 expression, which conferred more potent IE than stromal PD-L1 expression in murine CRC models (Kleinovink et al., 2017), can hence be a stable characteristic, similar to truncal IE through genetic mechanisms. Absent cancer cell PD-L1 expression in 15 cases was surprising in view of the abundant stromal PD-L1 positivity and the high mutation loads of all cases but this is consistent with previous reports in MMRd CRCs (Llosa et al., 2015).

The mechanisms determining cancer cell PD-L1 expression remain unknown and we correlated PD-L1 expression with immune cell infiltrates to investigate this. Because of the systematic bias towards higher immune infiltrates in lymph nodes, we only analysed primary tumor regions. Stromal PD-L1 expression correlated significantly with CD8 T-cells assessed by IHC (r=0.355, **Fig.3G**). Comparison with 15 immune cell subtypes inferred from RNA expression showed an even stronger correlation of stromal PD-L1 with activated CD8 T-cells (r=0.507), and also with activated CD4 and activated dendritic cells (**Fig.3H-I**). Thus, an adaptive immune response predominated by cytotoxic and helper T-cells strongly associated with stromal PD-L1 expression. In contrast, cancer cell PD-L1 expression did not significantly correlate with IHC quantified CD8 T-cells and only weakly with activated CD8 T-cells inferred from RNA expression (**Fig 3J**). Thus, we hypothesized that PD-L1 expression on cancer cells requires additional permissive factors.

We investigated whether any genetic drivers were specific to tumors with PD-L1 expressing cancer cells. *CD274*, which encodes PD-L1, was not amplified (Supplementary table 2). Truncal *FBXW7* driver aberrations were present in all 4 tumors with PD-L1+ cancer cells but also in 3 (2 truncal, 1 subclonal) that did not (**Fig.1D**). No other truncal drivers were enriched in tumors with PD-L1+ cancer cell. A previous report showed higher PD-L1 expression in CRCs with absent or low expression of the intestinal homeobox transcription factor CDX2 but this neither distinguished PD-L1 expression by stromal or cancer cells nor between MMRd and MMRp tumors (Inaguma et al., 2017). In our series, CDX2 RNA expression was significantly negatively correlated with PD-L1+ cancer cells (r=-0.683) but not with stromal PD-L1 (r=-0.269, **Fig 3K**). Remarkably, CDX2 expression was 24.7-fold lower in tumors with PD-L1+ cancer cells. We sought to validate this by IHC in an independent cohort of 23 MMRd CRCs (Supplementary table 6). Blinded to other data, a categorical scoring of CDX2 IHC staining intensity was performed as described (Dalerba et al., 2016). 17.4% of these tumors harbored PD-L1+ cancer cells. CDX2 staining was low (1+) or absent (0) in these whereas tumors without cancer cell PD-L1 expression predominantly showed strong (3+) or moderate (2+) CDX2 staining. This was statistically significant (**Fig.3L**). Moreover, analysis of RNA expression data from 57 colorectal cancer cell lines (Ghandi et al., 2019) confirmed a significant negative correlation of CDX2 and PD-L1 expression (**Fig.3M**).

Together, we found that stromal PD-L1 expression most strongly correlated with activated CD8 T-cells whereas expression by cancer cells was conditional on CDX2 expression loss.

### Co-evolution of genetic and immune landscapes

We next investigated how genetic IE mechanisms and the immune landscape co-evolved. Patterns of genetic IE evolution in cases where at least 3 primary tumor regions had been analysed classified these into three groups: 1. Tumors without evolution of genetic IE drivers, which we refer to as IE evolution stasis (T1, T2, T6, T13, T17, T18), 2. Tumors with subclonal IE driver evolution in some but not all regions (T4, T5, T14, T15), and 3. Tumors with pan-tumor IE through truncal aberrations or parallel evolution in all tumor regions (T3, T7, T12, T16, T19). We assessed whether the immune landscapes of these groups differed.

CD8 T-cell densities were significantly higher in group 2 (subclonal IE evolution) compared to groups 1 and 3 (**Fig.4A**). CD8 T-cell densities in group 1 (stasis) were similar to those in group 3 (pan-tumor IE evolution). Consistently, immune cell abundance inferred from RNA expression showed higher activated and effector memory CD8 T-cells, activated CD4 T-cells and immature B-cells in group 2 (**Fig.4B-C**). This was only significant if no multiple testing correction was applied, likely due to the small cohort size. Together, this indicates that the selection pressure from high CD8 T-cell infiltrates is the proximate cause for subclonal evolution in group 2. Conversely, the absence of this selection pressure likely explains the evolutionary stasis with respect to known IE mechanisms in group 1 whereas pan-tumor IE is likely responsible for the low CD8 T-cell infiltrates in group 3. Moreover, these subtypes explained the inter- and intra-tumor variability of CD8 T-cell infiltrates (**Fig.3B**).

**Figure 4:**
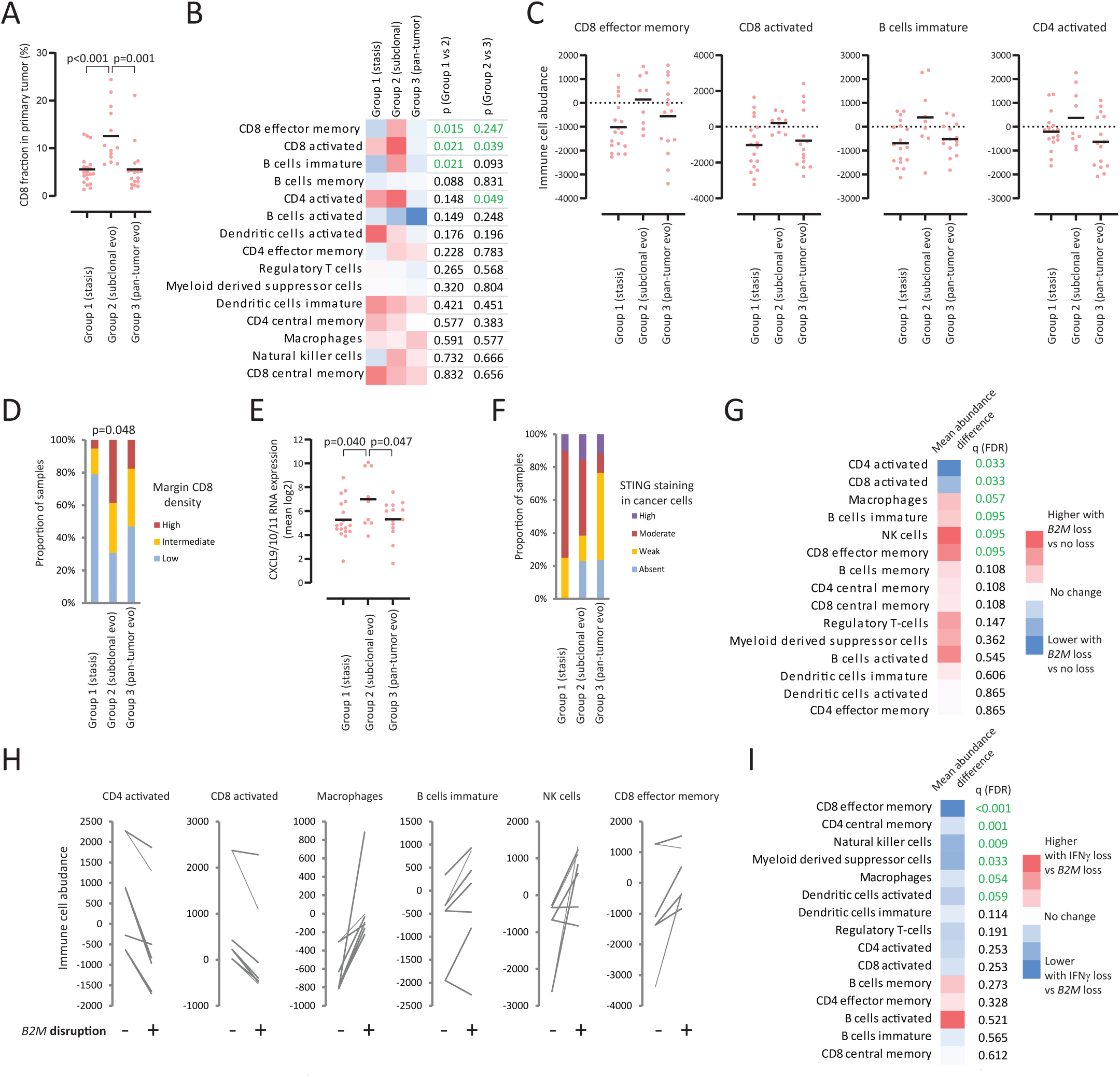
Co-evolution of genetic and immune landscapes. A. CD8 T-cell fraction (IHC) measured in 50 primary tumor regions from 15 MMRd CRCs that were classified into three groups based on IE driver evolution patterns. B. Abundance of 15 immune cell subtypes inferred from RNA expression data in the three evolution groups. C. Sample-level data and means (horizontal bars) for significant immune cell subtypes in panel B. D. CD8 T-cell density at the tumour margin by evolution group. E. Mean expression of the CD8 T-cell chemo-attractants CXCL9-CXCL11. F. Predominant staining intensity of STING (IHC) in cancer cells. G. Comparison of primary tumor regions with *B2M* defects (n=7) to all remaining primary tumor regions from the same cases (n=4). The heatmap shows the mean abundance in regions with *B2M* defects minus the mean abundance in regions without defects. H. Abundance difference of immune cell subtypes that were significant in G, comparing tumour regions with *B2M* disruption (+) to regions without disruption (-). I. Comparison of immune cell abundances in all primary tumor regions with defects of IFNγ pathway genes (n=7) to all remaining primary tumor regions with *B2M* disruption (n=15). The heatmap shows the mean abundance in regions with IFNγ defects minus the mean abundance in regions with *B2M* disruption. All horizontal bars show means. Significance analyses were performed with the Chi-squared test in panel D, the paired t-test in panel G, and the t-test in other panels.

We assessed why CD8 T-cell infiltrates were low in tumors with IE evolution stasis (group 1). This was not the consequence of lower mutation loads as they exceeded those in group 2 (median group 1: 56, median group 2: 51). Group 3 had the highest mutation load (median: 69). Assessing CD8 T-cell densities at the tumor margin by IHC (Supplementary table 2), we found that this was lowest in cases with evolutionary stasis (**Fig.4D**), suggesting that T-cell recruitment may be impaired. This was corroborated by a significantly lower mean expression of the CD8 T-cell chemo-attractants CXCL9-11 compared to group 2 (**Fig.4E**). Moreover, immuno-suppressive cell types such as myeloid derived suppressor cells, macrophages or regulatory T-cells were not enriched in group 1 (**Fig.4B**). The mean stromal PD-L1 score was highest in group 2 (1.92) followed by group 3 (1.41) and lowest in group 1 (1.25). Thus, PD-L1 induced T-cell apoptosis (Chiu et al., 2018) cannot explain low CD8 infiltrates in group 1. Activation of the cGAS-STING pathway may be necessary to promote immune recognition and CD8 T-cell infiltration in MMRd CRCs (Lu et al., 2021). Loss of STING expression occurs in CRCs (Xia et al., 2016) and may hinder immune recognition, yet we detected STING in all group 1 tumors (**Fig 4F**, Supplementary table 2). Overall our data show that impaired recruitment rather than intratumoral inactivation of CD8 T-cells explain the absence of IE evolution in group 1.

We next assessed how the evolution of specific IE drivers influences immune infiltrates within tumors. We compared immune profiles of primary tumor regions that lost genes in the HLA class I antigen presentation pathway (n=7) with proficient regions (n=4) in the same tumors. *B2M* was the affected gene in all cases. Activated CD4 and activated CD8 T-cells were significantly less abundant whereas macrophages, immature B-cells, NK-cells and CD8 effector memory cells were significantly higher in regions with defective *B2M* (**Fig.4G-H**). Inactivation of the IFNγ signalling pathway through mutations in *IFNGR1* or *JAK1* was a second common genetic IE mechanism. Their impact could not be assessed within tumors as only one case contained both subclonal driver and wild type regions. We therefore compared immune infiltrates between all primary tumor regions with IFNγ pathway inactivation (n=7) to the remaining ones with defective *B2M* (n=15). This revealed no significant difference in activated CD4 or CD8 T-cells but lower infiltrates for most other immune cell subtypes, including NK cells, demonstrating broad depression of immune cell infiltrates (**Fig.4I**). The enrichment of NK cells is an expected consequence of *B2M* loss and suggests opportunities for NK-cell therapies (Sade-Feldman et al., 2017). In contrast, the sparse infiltrates of most immune cell subtypes in tumors with IFNγ signalling defects shows that distinct treatment strategies are necessary for these.

### IE and metastatic dissemination

We finally explored whether IE mechanisms in cancer cells (including drivers in IE genes or cancer cell PD-L1 expression) associated with the presence of metastases. We found no difference between the three evolutionary subtypes (**Fig.5A**), and the proportion of tumors harboring any IE mechanism was similar between stage 1/2 tumors (50%) and stage 3/4 tumors (61.5%, **Fig.5B**). However, truncal IE were exclusively identified in stage 3/4 tumors (53.8%, **Fig.5C**), suggesting that early acquisition of these known IE mechanisms could increase the probability of metastasis development. This difference was statistically significant; nevertheless further studies are required to substantiate this in larger cohorts.

**Figure 5:**
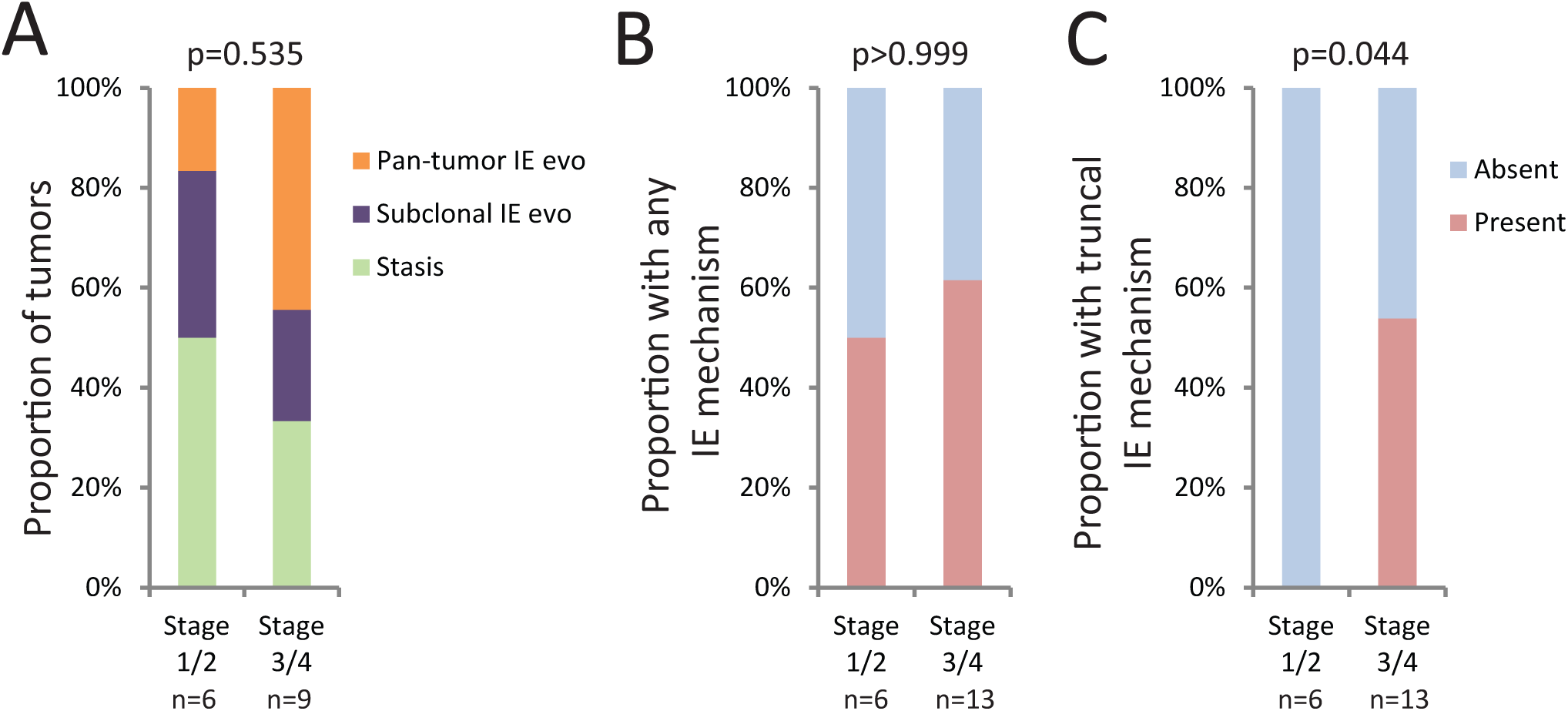
Immune evasion mechanisms in tumors with and without metastases. A. Three IE evolution subtypes by stage. B. Presence of any IE mechanisms in cancer cells (clonal or subclonal IE driver aberrations from Fig.2A or cancer cell PD-L1 expression) by stage. C. Presence of truncal IE mechanisms in cancer cells by stage. Cancer cell PD-L1 expression was considered equivalent to a truncal genetic IE driver as it was detected in all tumor regions where present. Statistical analyses were performed with the Chi-squared test for panel A and the Fisher’s exact test for panels B and C.

## DISCUSSION

MMRd CRCs have previously been subclassified by *BRAF* mutation status or MMR loss patterns but neither these nor mutation loads or heterogeneity metrics correlated with immune infiltrates in this cohort. Evolution analysis suggested a new taxonomy, with three subtypes, that provides an explanation for their distinct immune infiltrates: strong selection pressure from high CD8 T-cell infiltrates in tumors leads to subclonal IE; tumors that already established pan-tumor IE show low CD8 T-cell infiltrates; and tumors showing IE evolution stasis show low CD8 T-cells. Sparse CD8 T-cells at the tumor margin and low expression of T-cell chemo-attractants support impaired T-cell recruitment as the mode of immune escape in the latter.

All genetic IE evasion mechanisms analysed in these tumors have been associated with resistance to PD1 inhibitors (B2M, IFNγ pathway mutations) (Shin et al., 2017; Zaretsky et al., 2016) CTLA4 inhibitors (IFNγ pathway mutations) (Gao et al., 2016) or PD1/CTLA4 combination therapy (B2M) (Sade-Feldman et al., 2017) in lung cancers or melanoma. In contrast, a study in MMRd CRCs showed that some patients with reduced *B2M* protein expression did benefit from PD1/PD-L1 inhibition and that murine MMRd models with *B2M* inactivation were still sensitive to combined PD1/PD-L1 and CTLA4 treatment (Germano et al., 2021). ITH shown in our data is a major hindrance to accurately assess how AP or IFNγ pathway inactivation impacts immunotherapy responses in the clinic and if this differs depending on whether PD1 inhibition or combined PD1/CTLA4 blockade is used. ITH analysis of immune evasion drivers and subtype detection requires clonality analyses which are unlikely to be clinically feasible from tissue samples. However, circulating tumor DNA sequencing is effective in CRC and methods to determine driver aberration clonality have been developed (Knebel et al., 2020; Woolston et al., 2019). This can be readily implemented in clinical trials and may eventually allow stratification of patients to either PD1 inhibitors, more toxic CLTA4/PD1 combinations (e.g. for patients with *B2M* inactivation), or alternative therapies if neither of these are anticipated to be effective. Distinct IE drivers also had differing impact on immune infiltrates. Invigorating NK-cells could be a rational alternative treatment approach for tumors with truncal or subclonal inactivation of *B2M* (Sade-Feldman et al., 2017) whereas tumors with IFNγ signalling loss are unlikely to benefit based on our data.

Expression of PD-L1 by cancer cells was strongly associated with CDX2 loss, suggesting a permissive effect for PD-L1 upregulation. CDX2 is physiologically expressed in bowel epithelium where it controls intestinal differentiation and gene expression programs (Kaimaktchiev et al., 2004). Impaired CDX2 expression in cancers of the large and small bowel confers an increased recurrence risk (Dalerba et al., 2016; Jun et al., 2014; Tomasello et al., 2018) which has been thought to be a consequence of differentiation loss and increased invasiveness (Hryniuk et al., 2014). Our data indicates that an increased ability to suppress T-cell function through PD-L1 upregulation may contribute. In addition, cancer cell PD-L1 expression confers higher susceptibility to CPI treatment in murine models (Kleinovink et al., 2017; Lau et al., 2017). Whether CDX2 loss identifies tumors that particularly benefit from adjuvant CPIs should hence be investigated in ongoing trials (e.g. ATOMIC, ClinicalTrials.gov ID: NCT02912559). CDX2 is not known to play a role in immune regulation and our observation also raises questions regarding its function in infection or autoimmunity in the intestinal tract.

Finally, we revealed a hierarchy of driver aberration evolution in MMRd CRCs. This defines a model of MMRd CRC development where TGFBR family members, WNT and RTK/MAPK pathways are critical for cancer initiation whereas PI3-kinase, histone modifier genes, DDR genes and IE genes promote cancer progression. Inhibiting BRAF in CRCs with V600E mutations is well established in clinical practice (Kopetz et al., 2019). Our observation of truncal *ERBB2/ERBB3* driver mutations in some cases suggests opportunities to target these tumors with specific inhibitors.

## Acknowledgements

We thank the patients who donated samples for research and the clinical research teams for trial support and sample collection. The study was supported by funding to MG from the European Research Council under the European Union’s Horizon 2020 research and innovation program (grant agreement No. 820137) and from the Royal Marsden Hospital/ICR NIHR Biomedical Research Centre for Cancer. JR-B receives funding from a Río Hortega fellowship from the Institute of Health Carlos III (CM19/00087) and a 2020 TTD Research Grant from the Spanish Cooperative Group for the Treatment of Digestive Tumors (TTD). BRC receives funding from Cancer Research UK and The Wellcome Trust. The POLEM clinical trial was supported by a research grant from Merck KG. JM, JLA and JK received funding from the NIHR Imperial College Biomedical Research Centre. We thank Dr Gideon Coster at The Institute of Cancer Research, London for critical discussions.

## Author contributions

BRC and MG designed the study. BRC, MG, JR-B and JM obtained funding. BE, RE, SF, MH, DK, NM, LN, VP, MS, KS, TD, DC, IC, NSt and MG recruited patients to the POLEM trial. TD, DC, IC and NSt developed and coordinated the POLEM clinical trial. RB coordinated the clinical sample collection; and RC and AT co-ordinated clinical data collection. JR-B, JLA, JK, DL, GA and CF provided archival samples and data. BRC, HL-I, KvL, HBS and MD-A processed the tissue. BRC and LJB prepared sequencing libraries. IA and KF performed NGS sequencing. AW, BRC and MG performed bioinformatic analyses. BRC, TL, DB, NSi and HBS performed immunohistochemical staining. BRC, AW, MB, SG and MG interpreted the data. BRC, LJB and MG wrote the manuscript. All authors read and gave final approval of the manuscript

## METHODS

### Sample collection and preparation

Fourteen stage 3 MMRd CRCs were obtained from the UK POLEM phase 3 clinical trial (clinicaltrials.gov ID NCT03827044, ethics approval: 18/LO/0165)(Lau et al., 2020). The trial randomised patients with resected stage 3 MMRd CRC to either adjuvant chemotherapy followed by the PD-L1 inhibitor avelumab or to adjuvant chemotherapy alone but closed early and no outcome data was available. Five stage 1/2 and one stage 4 MMRd CRCs were obtained from the University Clinical Hospital of Santiago de Compostela, Spain (Galician Research Ethics Committee reference 2015/405). FFPE tissue (1 tumor block per case) for the validation cohort were obtained from UK POLEM clinical trial, from the University Clinical Hospital of Santiago de Compostela and from Imperial College Healthcare NHS Trust (Research Ethics Committee reference 14/EE0024). A pathologist-reported loss of MLH1, PMS2, MSH2 or MSH6 by immunohistochemistry was required for inclusion into this study. All patients had provided written informed consent for the use of tissues in research.

For multi-region analysis cases, FFPE blocks which represented the spatial extent of each resected tumor were selected (BRC or HL-I) and H&E stains were used by a pathologist (BRC) to identify tumor regions for manual macro-dissection from ten 10μm thick FFPE tissue sections per region. DNA and RNA were extracted with the Qiagen FFPE AllPrep kit according to manufacturer’s instructions; quantified and quality controlled using Qubit (Invitrogen), Tapestation and Bioanalyser (Agilent). Germline DNA was extracted from blood (n=19) or tumor adjacent non-malignant tissue (n=1).

### Targeted DNA sequencing

A targeted gene sequencing panel (Supplementary table 1) was designed to include all driver genes which are recurrently mutated in MMRd or MMRp CRCs or implicated in IE, if identified in at least two publications (Cortes-Ciriano et al., 2017; Gao et al., 2016; Giannakis et al., 2016; Grasso et al., 2018; von Loga et al., 2020; Zaretsky et al., 2016). Genes involved in antigen presentation pathways were also included despite identification in a single publication (Grasso et al., 2018), as this was a major area for investigation.

Sequencing libraries were prepared from tumor (target input 200 ng for DNA Integrity scores 3-8 or 500 ng for <3) and matched germline DNA (100 ng) using unique molecular identifiers for error correction and pooled according to the manufacturer’s protocol (Nonacus Cell3 Target). Paired-end 100 bp sequencing was performed by the Tumour Profiling Unit at the Institute of Cancer Research using an Illumina Novaseq with a target depth of 1000x in tumor regions and 100x in germline. Sequencing error correction and consensus BAM file preparation was performed using NonacusTools (v1.0). Samples were aligned to the ‘Homo_sapiens_assembly38’ reference files downloaded from the GATK resource bundle.

### Mutation calling

Mutect2 (v.4.1.4.1) was used to call somatic mutations. All samples belonging to the same tumor were analyzed simultaneously. The ‘af-only-gnomad.hg38.vcf.gz’ file was used as germline resource and an interval padding of 5 was specified. Resulting calls were annotated using annovar (v.2019-10-24) with build version hg38 and refgene, Cosmic v92 specified as protocol options. Calls were finally filtered for exonic/splicing using the ‘Func.refGene’ annotation. The primary calls were filtered using the following criteria: by variant allele frequency (VAF) >=5% and at least 5x higher VAF in at least one tumor region compared to the matched germline sample. Four metastases had low cancer cell content (T8 L, T9 L, T10 L, T12 L1) despite macro-dissection, likely reflecting the abundance of immune cells in these lymph node metastases. For these samples, mutations were accepted as present if they had a VAF >=1%. Median VAFs of all ubiquitous mutations per case were calculated for each region and 2x the median VAF of ubiquitous mutations was used to calculate the cancer cell content of each tumor region, which is a reliable estimate in near diploid tumors such as MMRd CRC.

### HLA mutation analysis

The algorithms used in the HLA mutation calling pipeline have been developed for hg19 and so all samples were first realigned using NonacusTools and the ‘Homo_sapiens_assembly19’ reference from the GATK resource bundle. POLYSOLVER(Shukla et al., 2015) v4 was used for mutation detection in HLA genes. The docker image was converted to singularity format and run using the singularity container (v3.6.4). A minor adjustment was necessary to correct for a missing ‘SAMTOOLS_DIR’ definition in the primary scripts. HLA typing was run on all germline samples using the ‘shell_call_hla_type’, with parameters race = Unknown, includeFreq = 1 and insertCalc = 0. Mutations in HLA genes were first identified using the ‘shell_call_hla_mutations_from_type’ script and were subsequently annotated using the ‘shell_annotate_hla_mutations’ script. HLA disrupting SNV and INDEL calls that met the criteria of VAF >5% were manually reviewed for confirmation of affected tumour regions using IGV(Robinson et al., 2011).

### DNA copy number analysis

CNVkit (v.0.9.8) was used for DNA copy number analysis. The *autobin* function suggested an antitarget bin size of 225585 based on sample depth. CNVkit was then run in paired batch mode using the hybrid protocol, antitarget bin size 225585, minimum antitarget bin size of 500 and --drop-low-coverage set to active. Absolute gene level copy number estimates for genes on autosomes (excluding polymorphic HLAs) were then generated using the .cnr file. Log2 values were adjusted for tumor purity and a weighted mean calculated across bins corresponding to the same gene. This process was repeated for the .cns file to generate segmented absolute gene level estimates and values were rounded to integer copy number. The tumor content of five lymph node metastases was too low to generate accurate copy number data (median truncal VAF<10%). Copy number information from the available primary tumor sample (T8, T9, T10) and in case of T18 L1 from the LN metastasis had a higher tumor content (median truncal VAF>10%) and was located on the same clade (T18 L2) and for T12 L1 the copy number data from the most closely related primary tumor region (T12 R2) was used. This is appropriate for MMRd tumors as copy number variation is limited.

### Identification of likely driver aberrations

Distinct criteria were used to define likely driver aberration in oncogenes vs. tumor suppressor and IE genes. The list of genes that were assessed in each group (Supplementary table 3) was selected based on evidence of driver function and classification as tumor suppressor, oncogene or relevance in IE in the COSMIC cancer gene census or in relevant publications.

#### Criteria for the identification of likely drivers in oncogenes

Any hotspot mutation, defined as mutations leading to amino acid changes that have been identified at least in 10 tumors in v92 of the COSMIC cancer mutation database, in an oncogene of interest was considered a likely driver.

#### Criteria for the identification of likely drivers in tumor suppressor genes and IE genes

These genes usually only have driver function when all alleles in the genome of the cancer cell are inactivated. We evaluated mutations that disrupt gene function (frameshift mutations, premature stop codons, splice-site mutations and mutations that disrupt the signaling peptide in B2M) as well as gene copy number to determine driver status. We used the copy number of each gene, the median ubiquitous VAF (as a measure of cancer cell content in a sample) and the VAF of disrupting mutations to calculate the number of gene copies carrying a mutation and then compared this to the number of total gene copies. This was done for each tumor region or metastasis individually. As indel mutations in MMRd tumors preferentially occur in repetitive DNA sequences which are unstable, more than one nucleotide can be lost or gained in a sequential fashion. We therefore combined the VAF of multiallelic indel mutation calls if those can arise from sequential deletion or from sequential insertion events (e.g. T1 *TGFBR2* chr3:30650379: VAFs of GAA>GA and GAA>G deletions were combined).

The following formulas, derived from(Woolston et al., 2019), were used to calculate the number of copies mutated in each samples:

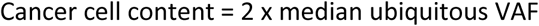

Expected clonal mutation VAF if one of a total of n gene copies per cancer cell is mutated:

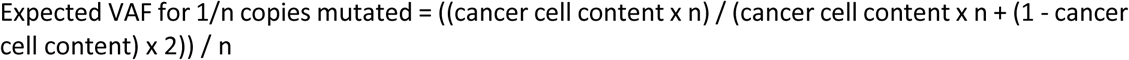

Simplified to:

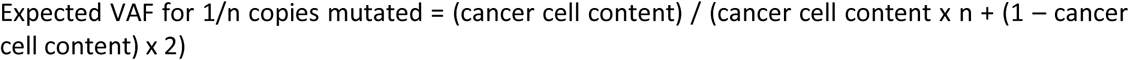

In the final step we divide the observed VAF for each mutation by the expected VAF when one of n copies is mutated to obtain the number of DNA copies mutated:

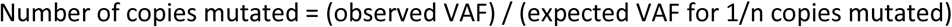

Biallelic inactivation can occur through one of the following:

1. A disrupting mutation (dm) combined with loss of heterozygosity (LOH)/loss of wild type alleles.
2. Two disrupting mutations (dm/dm) of which at least one is clonal in a tumor region so that the second disrupting mutation has to be nested within the same subclone.
3. Loss of all copies of a gene, ie resulting in copy number 0.

The following criteria that allowed up to 25% lower copy number of a mutation than the exact clonal estimate (to account for measurement inaccuracies) were used to define these instances:

##### 1. dm/LOH

- gene copy number 1 and a disrupting mutation with any number of copies mutated
- gene copy number 2 and 1.5 or more copies harboring a disrupting mutation
- gene copy number 3 and 2.3 or more copies harboring a disrupting mutation

If these criteria were fulfilled, additional disrupting mutations were ignored as functionally irrelevant because the gene was already inactivated through the first dm and LOH. The only exception were cases where a dm/LOH pattern was in conflict with the phylogenetic tree structure where we also tested whether dm/dm patterns showed a better fit and led to lower heterogeneity.

##### 2. dm/dm

Two independent disrupting mutations can either be present together in the same cancer cell clone or segregate in different subclones. As only the former can lead to biallelic inactivation, we established the following criteria to identify this. These are based on the principle that if one mutation is clonal then the second mutations has to be nested within the same clone.

- gene copy number 2 and at least 0.8 copies mutated by one disrupting mutation and any number of copies by the second one
- gene copy number 3 and at least 1.5 copies mutated by one disrupting mutation and any number of copies by the second one Importantly hotspot mutations in tumor suppressor or IE genes were considered equal to a disrupting mutation as recurrent acquisition indicates that they also likely impair gene function. As described in the results, we assessed whether two independent dm in *B2M* were present on different alleles which is a further requirement for inactivation. For this, BAM files were loaded into the Integrative Genomics Viewer IGV(Robinson et al., 2011), paired end reads were displayed and both mutations were visualized.

#### Exceptions to the above rules for specific genes

Most hotspot mutations in *TP53* have dominant negative effects. We therefore considered a single *TP53* hotspot mutation without LOH as a likely driver. *FBXW7* shows haploinsufficiency as heterozygous disrupting mutations or hotspot mutations can establish loss of function(Yeh et al., 2018). Thus, a single disrupting or hotspot mutation even without LOH was considered a driver. Kinase dead mutation in the tumor suppressor gene *PTEN* have been shown to have a dominant negative effect and these were also considered drivers regardless of LOH(Leslie and Longy, 2016; Smith and Briggs, 2016). For *B2M* we also considered mutations of the signal peptide, which is required for import into the endoplasmic reticulum. The N-terminal signal peptide location in *B2M* was retrieved with SignalP-5.0 (http://www.cbs.dtu.dk/services/SignalP/, Supplementary figure 4). One tumor showed a mutation (L13R) establishing a positively charged arginine that disrupts the hydrophobic signal peptide, likely precluding ER import.

#### Exceptions made because of phylogenetic tree conflicts

Out of 191 different driver aberrations that were identified, 2 exceptions were made from the above rules as the results generated phylogenetic conflicts that triggered a reassessment of VAF and copy number data which provided an alternative solution which was consistent with the trees. T6: *APC* was inactivated by copy number loss to one copy and one disrupting mutation in R1 and R3 but showed two copies and a disrupting mutation of one copy in R2. This would have required acquisition on two separate branches of the tree and was unusual as all other *APC* driver aberrations were truncal. We assessed copy number plots and found a narrow deletion at the APC position in all regions that was missed in T2 through under-segmentation. We therefore used unsegmented gene-level copy number data which showed one *APC* copy in R2. T13: *CIC* had 3 copies in L1 which, with only two mutated copies, would make this the only region without biallelic loss which is difficult to explain based on the phylogenetic tree. Identical to the approach above, we used unsegmented gene-level copy number data which showed two copies in L1, indicative of truncal acquisition.

All likely drivers identified are summarized with the supporting evidence in Supplementary table 4.

### Pylogenetic tree reconstruction

Mutation calls were transformed into a binary presence/absence matrix for all tumor regions per case. These were analysed with the PHYLIP Pars algorithm in the T-Rex online analysis suite(Boc et al., 2012) using a string of zeros to represent the germline as the outgroup and default settings. For Figure 1D phylogenetic trees were redrawn with branch lengths corresponding to the Pars output. Identified drivers were added to the tree at the branch where they had been acquired. Phylogenetic conflicts occurred for three driver aberrations whose regional distribution violated the tree structure. Each of these was considered to have occurred through multiple independent events (*AIRD1A* in T17, *RNF43* in T18, *BLM* in T7).

### Immunohistochemistry

3μm thick FFPE sections were stained for CD8 (1:20, C8/144B, Dako) on a Roche Ventana autostainer. Slides were scanned at 20x objective magnification and tumor regions correlating to the macro-dissected regions were manually annotated by a pathologist (BRC) using Qupath 0.2.3. Areas of tissue damage, necrosis and the stromal area of the tumor margin were excluded, so that the viable intra-tumour cancer cell and stroma areas were represented. The CD8+ fraction was calculated as a percentage of all nuclei detected with Qupath 0.2.3 in the annotated region applying a Nucleus:DAB OD mean threshold of 0.3 to define CD8 positive cells:

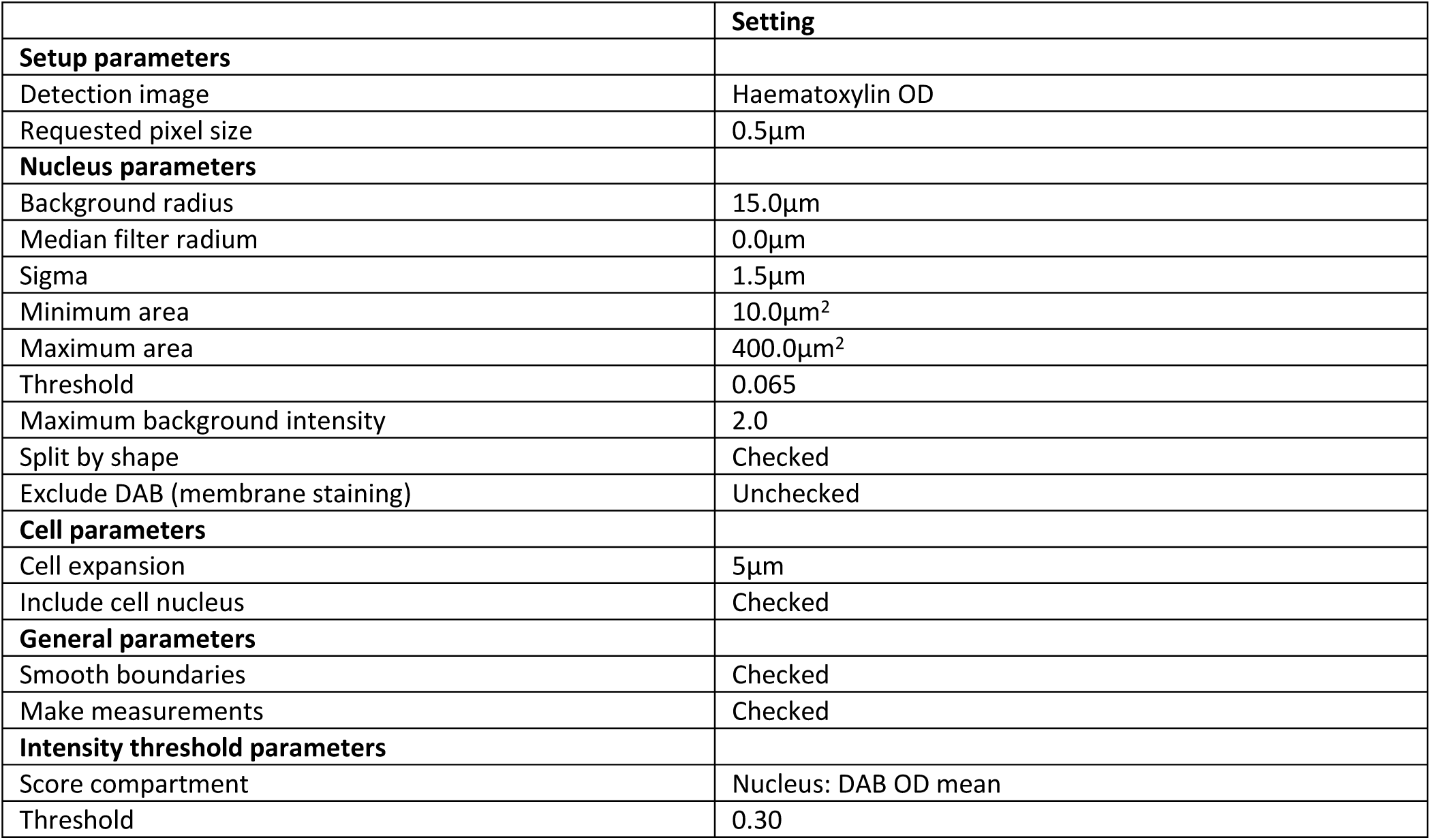

The density of CD8+ immune cells at the stromal margin of the tumor was then assessed by a pathologist (BRC) and categorized as low (absent to infrequent CD8 positive cells), moderate (intermediate CD8 positive cell density) or high (marked CD8 positive cell density).

STING (1:150, D2P2F, Cell Signaling Technology) and PD-L1 (1:100, E1L3N, Cell Signaling Technology) immunohistochemistry were performed on Leica Bond autostainer. Slides were scanned at x20 objective magnification and the tumor region, as defined at the time of the CD8 IHC analysis, was manually scored by a pathologist (BRC) blinded to other data. PD-L1 staining could be identified within the cytoplasmic and membranous compartments of stromal and tumor cells. Complete membranous staining was defined as positive and the proportion of positive stromal cells and cancer cells were each categorized with accepted cut-offs: 0% (negative), 1-4% (1+ positive), 5-49% (2+ positive) and 50%+ (3+ positive) (Moller et al., 2021). STING staining was identifiable within the cytoplasmic compartment of all normal epithelial and stromal cell types of colonic tissue. Cancer cell STING staining intensity was categorized as absent (0), weak (1+), moderate (2+) and strong (3+).

IHC for the validation cohort was performed on consecutive sections for each case. PD-L1 IHC was performed and the proportion of cancer cells positive was assessed as described above, blinded to other data. CDX2 IHC (1:100, EPR2764Y) was performed on a Leica Bond autostainer. CDX2 staining was identifiable within the nuclear compartment of normal colonic epithelial cells and on normal appendix epithelial cells which had been added as positive controls. The cancer cell nuclear intensity of CDX2 staining was categorized into 4 groups, absent (0), weak (1+), moderate (2+) and strong (3+) as described previously in a large cohort of CRCs (Dalerba et al., 2016) and blinded to other data.

### 3’-RNA sequencing

3’-RNA sequencing libraries were generated from all 71 samples. Unique dual indexed sequencing libraries were prepared from 500ng of RNA using Lexogen QuantSeq 3’mRNA-Seq FWD library prep kit and pooled according to the manufacturer’s protocols. Single-end 100 bp sequencing was performed by the Tumour Profiling Unit at the Institute of Cancer Research using an Illumina Novaseq. Resulting fastq were uploaded to the BlueBee Genomics platform for analysis. Briefly, this pipeline involves STAR alignment, HTSeq read counting and DESeq2 normalization. Nine samples failed QC due to fewer than 10000 genes detected, leaving 62 samples for analysis.

### Immune cell abundance analysis using ssGSEA

ssGSEA was performed with the updated gene symbols from(Charoentong et al., 2017) for 15 immune cell types. Normalized RNA sequencing data from 60 tumor samples (excluding T20 which was identified as an MMRp tumor) was analysed with these signatures using the ssGSEA module on the GenePattern workbench (https://cloud.genepattern.org) with default settings. For each cell type, the median of all samples was subtracted from the abundance values of individual samples.

### Expression analysis of colorectal cancer cells lines

mRNA expression data for PD-L1/CD274 and CDX2 were downloaded from the Cancer Cell Line Encyclopedia dataset (Ghandi et al., 2019) on the cBIO portal (www.cbioportal.org). Expression values were offset by 1 and log2 transformed for linear regression analysis.

### Statistical analyses

Statistical analyses were performed with the GraphPad PRISM. All p-values are two tailed and p<0.05 was considered significant. Where appropriate, multiple testing correction was performed with the False Discover Rate method by Benjamin, Krieger and Yekutieli and a q-value of <0.1 was considered significant. Hierarchical Euclidian clustering for the heatmap in Figure 3D was performed with the Morpheus clustering software (https://software.broadinstitute.org/morpheus/) and the inbuilt t-test and multiple testing correction were used. For the immune cell comparison of *B2M* loss within tumors, RNA sequencing data for only one region without *B2M* loss was available for 3 tumors (T3, T5, T14). This was used as the comparator for each of the regions with *B2M* loss and the paired t-test was applied. Normalized RNA expression values were offset by 1 and Log2 transformed before statistical significance analysis.

