## Supplementary figures 1,2, 3 and 4. Supplementary tables 1, 3, 5 and 6 for "Genetic and immune landscape evolution defines subtypes of MMR deficient colorectal cancer"

Challoner et al.

**Supplementary figure 1:** Linear regression model for extrapolation to exome **Page 2**

**Supplementary figure 2**: Ubiquitous mutation load by *BRAF*, MMR pattern and stage **Page 2**

**Supplementary figure 3**: Correlation of CD8 T-cell fractions by IHC vs immune cells inferred from RNA expression **Page 3**

**Supplementary figure 4**: Signal peptide location in the B2M protein sequence **Page 3**

**Supplementary table 1**: 194 genes included in the targeted sequencing panel **Page 4**

**Supplementary table 2**: Patient characteristics, sequencing metrics, IHC data and ssGSEA immune cell abundance **Excel file**

**Supplementary table 3**: Genes analysed for driver aberrations **Page 5**

**Supplementary table 4**: Data supporting the identified driver aberrations **Excel file**

**Supplementary table 5**: Correlation of CD8 T-cell fractions with mutation loads **Page 5**

**Supplementary table 6:** Patient and pathological characteristics of the validation cohort **Page 6**

**Supplementary figure 1**: Linear regression model generated with GraphPad PRISM by analysing non-silent mutation calls in the exome vs those in 191 target genes (excluding class I HLA genes) in our targeted sequencing panel for 518 MSI and MSS CRCs (POLE cases were excluded). TCGA pan-cancer whole-exome mutation data was downloaded from the cBio portal. The equation for the linear regression line and the $r^2$ value generated by PRISM are shown.

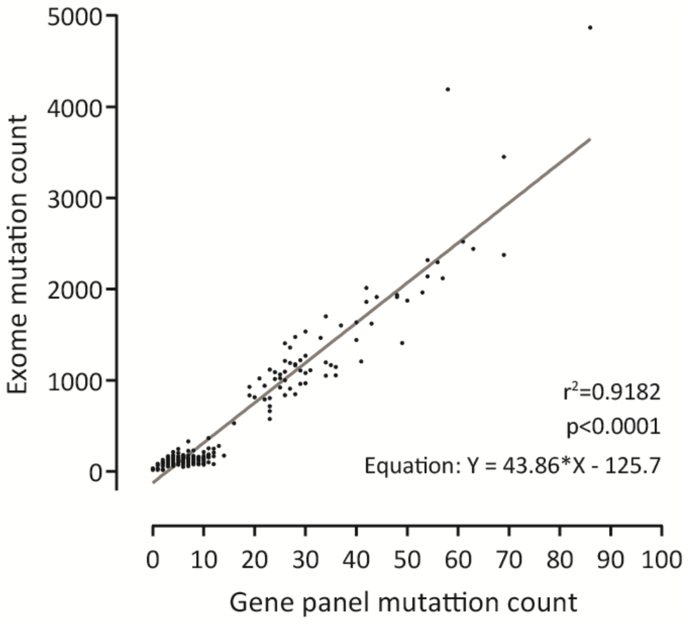


**Supplementary figure 2:** Ubiquitous mutation load by BRAF, MMR loss pattern and stage, horizontal bars indicate medians. The Mann-Whitney test was used to assess significance.

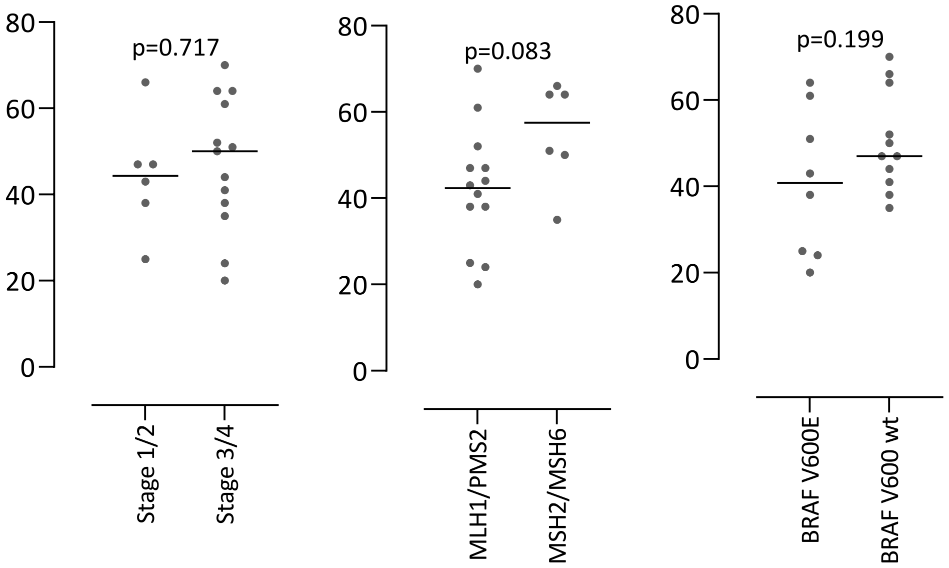

**Supplementary figure 3:** Pearson correlation analysis of CD8 T-cell fractions assessed by IHC vs immune cell subtype abundances assessed by ssGSEA in 60 tumor regions from which RNA-sequencing data was available. The Pearson correlation coefficient and p values were calculated and multiple testing correction using the false discovery rate (FDR) approach was applied to generate q values. A q value <0.1 was considered significant.

| | r | q (FDR) |
|---|---|---|
| CD8 activated | 0.525 | <0.001 |
| CD4 activated | 0.358 | 0.024 |
| CD8 effector memory | 0.347 | 0.024 |
| Myeloid derived suppressor cells | 0.344 | 0.024 |
| B cells immature | 0.338 | 0.024 |
| B cells activated | 0.288 | 0.061 |
| Dendritic cells activated | 0.274 | 0.069 |
| CD8 central memory | 0.248 | 0.097 |
| Dendritic cells immature | 0.238 | 0.097 |
| Regulatory T cells | 0.237 | 0.097 |
| CD4 effector memory | 0.203 | 0.154 |
| Macrophages | 0.194 | 0.163 |
| Natural killer cells | 0.102 | 0.452 |
| CD4 central memory | 0.100 | 0.452 |
| B cells memory | 0.044 | 0.696 |

**Supplementary figure 4:** Signal peptide location in the B2M protein sequence.

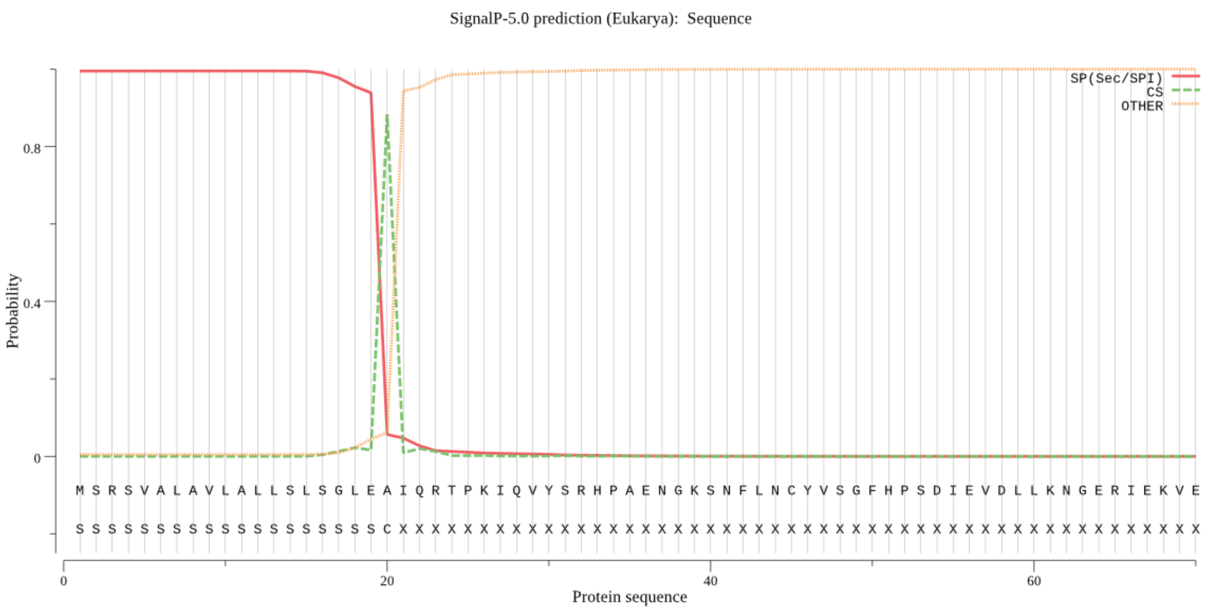

**Supplementary table 1:** 194 genes included in the targeted sequencing panel.

| | | | | | | |
|---|---|---|---|---|---|---|
| ACVR1B | BRCA2 | ERAP2 | KMT2D | NLRC5 | RIPK2 | TCF7 |
| ACVR2A | CALR | ERBB2 | KRAS | NRAS | RNF128 | TCF7L2 |
| ADAM30 | CANX | ERBB3 | KRTAP4-5 | ORC2 | RNF43 | TGFBR1 |
| AKAP7 | CASD1 | FBXW7 | LARP4B | PA2G4 | RNF6 | TGFBR2 |
| AKT1 | CASP8 | FGFR1 | MAP2K1 | PAX6 | RNGTT | TGIF1 |
| ALX4 | CD274 | FGFR2 | MAP7 | PBRM1 | RPL22 | TM9SF3 |
| AMER1 | CDC27 | FHOD3 | MBD6 | PDCD1LG2 | RPRD1A | TNFRSF4 |
| ANKMY2 | CDH1 | FRMD4A | MET | PDIA3 | RPS27L | TNFRSF9 |
| ANKRD46 | CDKN2A | FZD3 | MIER3 | PHACTR1 | RWDD4 | TP53 |
| APC | CENPH | HLA-A | MITF | PHGR1 | SALL4 | TRAM1L1 |
| ARHGAP5 | CIC | HLA-B | MLH1 | PIK3CA | SDHA | TRIM48 |
| ARID1A | CIITA | HLA-C | MLH3 | PIK3R1 | SDHAF2 | TRIM51 |
| ARID1B | CPEB2 | HNRNPA2B1 | MSH2 | PLEKHA6 | SLC25A36 | UBQLN2 |
| ARID2 | CREBBP | HSPA5 | MSH3 | PMS1 | SLC3A2 | UBR2 |
| ASXL1 | CRTC1 | IDH1 | MSH6 | PMS2 | SMAD2 | UBR5 |
| ATF5 | CTCF | IDH2 | MT3 | POLD1 | SMAD3 | VPS35 |
| ATM | CTNNB1 | IFNGR1 | MTOR | POLE | SMAD4 | WNK4 |
| ATP6V1B1 | CUL5 | IFNGR2 | MVK | PRDM2 | SMARCA4 | WNT1 |
| ATR | DHX40 | ING1 | MYC | PRKDC | SMCP | WNT16 |
| ATXN2L | DIAPH1 | ITGA2 | MYO1B | PTEN | SNAPC1 | WWOX |
| AXIN2 | DICER1 | JAK1 | MYOCD | PTH2 | SOX9 | XYLT2 |
| B2M | DNAH5 | JAK2 | NBN | PTPN12 | SPRR1A | YWHAE |
| BARD1 | DOCK3 | JUN | NDUFA10 | QKI | STAT1 | ZBTB20 |
| BCL9L | DUSP16 | KCTD20 | NDUFAF6 | RAD17 | TAP1 | ZDHHC8 |
| BLM | EDNRB | KLF3 | NEFH | RB1 | TAP2 | ZFP36L2 |
| BMPR2 | EGFR | KLF5 | NF1 | RBM10 | TAPBP | ZNRF3 |
| BRAF | EMILIN1 | KLHL42 | NF2 | RFX5 | TCERG1 | |
| BRCA1 | ERAP1 | KMT2C | NGRN | RGMB | TCF20 | |

**Supplementary table 3:** Known oncogenes, tumor suppressor genes and immune evasion genes (involved in HLA-I antigen presentation or IFNγ signalling) which were assessed for driver aberrations.

| Oncogene | Tumor suppressor | | | Immune evasion |
|---|---|---|---|---|
| AKT1 | ACVR1B | BRCA2 | PMS2 | B2M |
| BRAF | ACVR2A | CDH1 | POLD1 | CALR |
| CTNNB1 | AMER1 | CDKN2A | POLE | CANX |
| EGFR | APC | CIC | PRDM2 | ERAP1 |
| ERBB2 | ARID1A | FBXW7 | PTEN | ERAP2 |
| ERBB3 | ARID1B | KMT2C | RNF43 | IFNGR1 |
| FGFR1 | ARID2 | KMT2D | RPL22 | IFNGR2 |
| FGFR2 | ASXL1 | MLH1 | SMAD2 | JAK1 |
| KRAS | ATM | MSH2 | SMAD3 | JAK2 |
| MAP2K1 | ATR | MSH3 | SMAD4 | NLRC5 |
| MET | AXIN2 | MSH6 | SMARCA4 | PDIA3 |
| MTOR | BCL9L | NF1 | TCF7L2 | STAT1 |
| NRAS | BLM | NF2 | TGFBR2 | TAP1 |
| PIK3CA | BMPR2 | PBRM1 | TP53 | TAP2 |
| | BRCA1 | PIK3R1 | | TAPBP |

**Supplementary table 5:** Spearman correlation coefficient (r) of mean CD8 fraction in primary tumor regions vs. mutation metrics in 19 MMRd CRCs and significance analysis.

| | Mean CD8 fraction vs. | | | |
|---|---|---|---|---|
| | mean mutation load | mean heterogeneous mutation fraction | ubiquitous mutations | ubiquitous indels |
| Spearman r | -0.2442 | -0.3982 | -0.1148 | -0.00527 |
| p | 0.3137 | 0.0913 | 0.6397 | 0.9829 |

**Supplementary table 6:** Patient and pathological characteristics of the validation cohort of 23 MMRd CRCs.

| | MMRd CRC Cohort (n=23) |
|---|---|
| **Median age at resection (range)*** | 67.7 (36.8-79.7) |
| **Sex** | |
| Male | 17% (4) |
| Female | 44% (10) |
| Unknown | 39% (9) |
| **Stage (AJCC/UICC 8th edition)** | |
| 1 | 4% (1) |
| 2 | 22% (5) |
| 3 | 70% (16) |
| 4 | 4% (1) |
| **Predominant Differentiation** | |
| Well to moderate | 39% (9) |
| Poor | 39% (9) |
| Mucinous | 22% (5) |

*status known (n=14)
